# Proton Relaxometry of Tree Leaves at Hypogeomagnetic Fields

**DOI:** 10.1101/2023.12.08.570690

**Authors:** Anne M. Fabricant, Piotr Put, Danila A. Barskiy

## Abstract

We report on a cross-species proton-relaxometry study in *ex vivo* tree leaves using nuclear magnetic resonance (NMR) at 7 μT. Apart from the intrinsic interest of probing nuclear-spin relaxation in biological tissues at magnetic fields below Earth field, our setup enables comparative analysis of plant water dynamics without the use of expensive commercial spectrometers. In this work, we focus on leaves from common Eurasian evergreen and deciduous tree families: Pinaceae (pine, spruce), Taxaceae (yew), Betulaceae (hazel), Prunus (cherry), and Fagaceae (beech, oak). Using a nondestructive protocol, we measure their effective proton *T*_2_ relaxation times as well as track the evolution of water content associated with leaf dehydration. Newly developed “gradiometric quadrature” detection and data-processing techniques are applied in order to increase the signal-to-noise ratio (SNR) of the relatively weak measured signals. We find that while measured relaxation times do not vary significantly among tree genera, they tend to increase as leaves dehydrate. Such experimental modalities may have particular relevance for future drought-stress research in ecology, agriculture, and space exploration.

## 1 INTRODUCTION

The essential problem of measuring water content and dynamics in plants may seem simple enough. To this day, however, the water-monitoring toolbox remains surprisingly limited, especially where nondestructive techniques are concerned. A standard approach involves desiccating harvested plant organs and comparing their fresh and dry weights [1, 2, 3, 4]. In recent years, proton (^1^H) nuclear magnetic resonance (NMR) has emerged as a promising alternative technique, due to its sensitivity to water protons. ^1^H NMR has found a number of plant-related applications—including measurement of water content in lumber wood [5], investigation of moisture stress in agricultural seeds [6, 7], and characterization of microbial interactions in soil [8]. Techniques based on proton relaxometry are now particularly relevant within food science [9, 10, 11], where magnetic resonance imaging (MRI) is also employed [12]. However, commercially available NMR spectrometers typically do not have suitable geometries for measurement of intact plants or plant organs.

Leaves are arguably the most critical actor in the plant water cycle, given that a majority of transpiration and systemic water loss occurs there. Despite this, the use of proton NMR in leaf water studies is far from mainstream, although some relaxometry work has been carried out with low-field benchtop spectrometers. Notably, relaxometry of senescing rapeseed leaf pieces (excised discs) at 20 MHz was investigated using a Carr-Purcell-Meiboom-Gill (CPMG) protocol, indicating an increase in some *T*_2_ (spin-spin relaxation time, also known as coherence time) components in older leaves [13, 14]. A similar study at 20 MHz demonstrated the utility of *T*_2_ relaxation for phenotyping and detection of water stress in excised leaves of young potted tobacco plants [15]. At high field, proton *T*_2_ relaxometry was applied to structural water studies in maple leaves [16]. In addition, high-field solid-state proton NMR was shown to be effective for studying relaxation properties of dried leaves and leaf litter even when little water is present, by revealing the molecular fingerprint of plant metabolites and biopolymers [17].

The flexibility and portability of low-field NMR—loosely defined as corresponding to magnetic fields ranging from Earth field up to a few tesla, above which superconducting or hybrid superconducting/electromagnets would be required—also offers potential for taking devices directly into the field, forest, or greenhouse. One major example is the realization of *in vivo* and *ex vivo* water-proton relaxometry of intact leaves from potted agricultural plants, as well as wild shrubs and oak and poplar trees, using a unilateral 18 MHz spectrometer [18]. Currently, noncommercial low-field relaxometers are being developed which enable portable *in vivo* measurement of even larger plant leaves and organs [19]. Such devices complement other novel non-NMR modalities for nondestructive monitoring of leaf water potentials, e.g. nanobiosensors [20]. In trees, water transport in living tree trunks and branches has been studied using custom low-field MRI and NMR devices [21, 22].

In traditional NMR systems based on inductive detection, the tradeoff between portability and achievable signal-to-noise ratio (SNR) limits how low of a magnetic field can be reasonably used for measurement of intact biological systems, where signal strengths tend to be relatively weak. It has been shown theoretically that below proton resonance frequencies of around 50 MHz, detection using atomic (optically pumped) magnetometers can offer better intrinsic sensitivity than that attainable with inductive pickup coils [23]. The atomic-magnetometry detection modality has been instrumental in the subfield of zero-to-ultralow-field (ZULF) NMR [24, 25, 26], where ULF is commonly used in literature to refer to fields below the geomagnetic (Earth) field of tens of microtesla, such that magnetic shielding or active field cancellation is required. Due to varying definitions of ULF by different authors, some absolute [27] and others referenced to the spin system under study—e.g., *J*-coupling between spins dominates Zeeman interactions with the external field [28]—we choose instead to use the unambiguous term “hypogeomagnetic” in this publication. The hypogeomagnetic regime has already been used for direct detection of biomagnetic fields produced by plant electrical activity, including action potentials and wounding potentials [29, 30, 31]; however, according to our understanding, NMR signals originating from plants have not yet been explored in this regime.

In addition to the fundamental question of how proton relaxation properties behave at hypogeomagnetic fields, the regime is interesting from a practical NMR standpoint, due to the low cost, portability, and low energy consumption of experimental components. Although NMR detection using superconducting-quantum-interference-device (SQUID) magnetometers [27, 32] offers comparable sensitivity to atomic magnetometers at hypogeomagnetic fields (as well as a larger frequency bandwidth), the need for bulky cryogenic cooling limits the applicability of SQUID-based devices. The smaller footprint of atomic magnetometers also allows placement of multiple sensors around a sample, rather than in a single detection plane.

To our knowledge, the work reported here represents the broadest cross-species NMR-relaxation study of tree leaves at any magnetic field, and the first to incorporate both evergreen and deciduous varieties. Through systematic nondestructive measurement of intact leaves from seven different tree genera—spruce, pine, yew, hazel, cherry, beech, and oak—we endeavored to investigate variation in water-proton signals and relaxation times among genera. In targeted studies of spruce and oak samples, we also sought to track the evolution of these parameters as a function of leaf dehydration. While relative proton signal correlates with wet mass upon dehydration, *T*_2_ times tend to increase, indicating the possible presence of compartmentalized water reservoirs with higher water mobility surviving upon dehydration.

## 2 MATERIALS AND METHODS

### 2.1 Relaxometry setup

Our gradiometric quadrature detection scheme is shown in Figure 1A, where two commercially available dual-axis vector magnetometers [33] are placed orthogonally in the *x*-*y* plane. If the sensors, denoted 1 and 2, are positioned symmetrically around a magnetic dipole initialized along 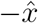 and undergoing Larmor precession clockwise in the plane at positive angular frequency *ω*_0_, we can write the following (ideal) expressions for the oscillating magnetic field sensed by the four magnetometer channels as a function of time *t*:

**Figure 1.**
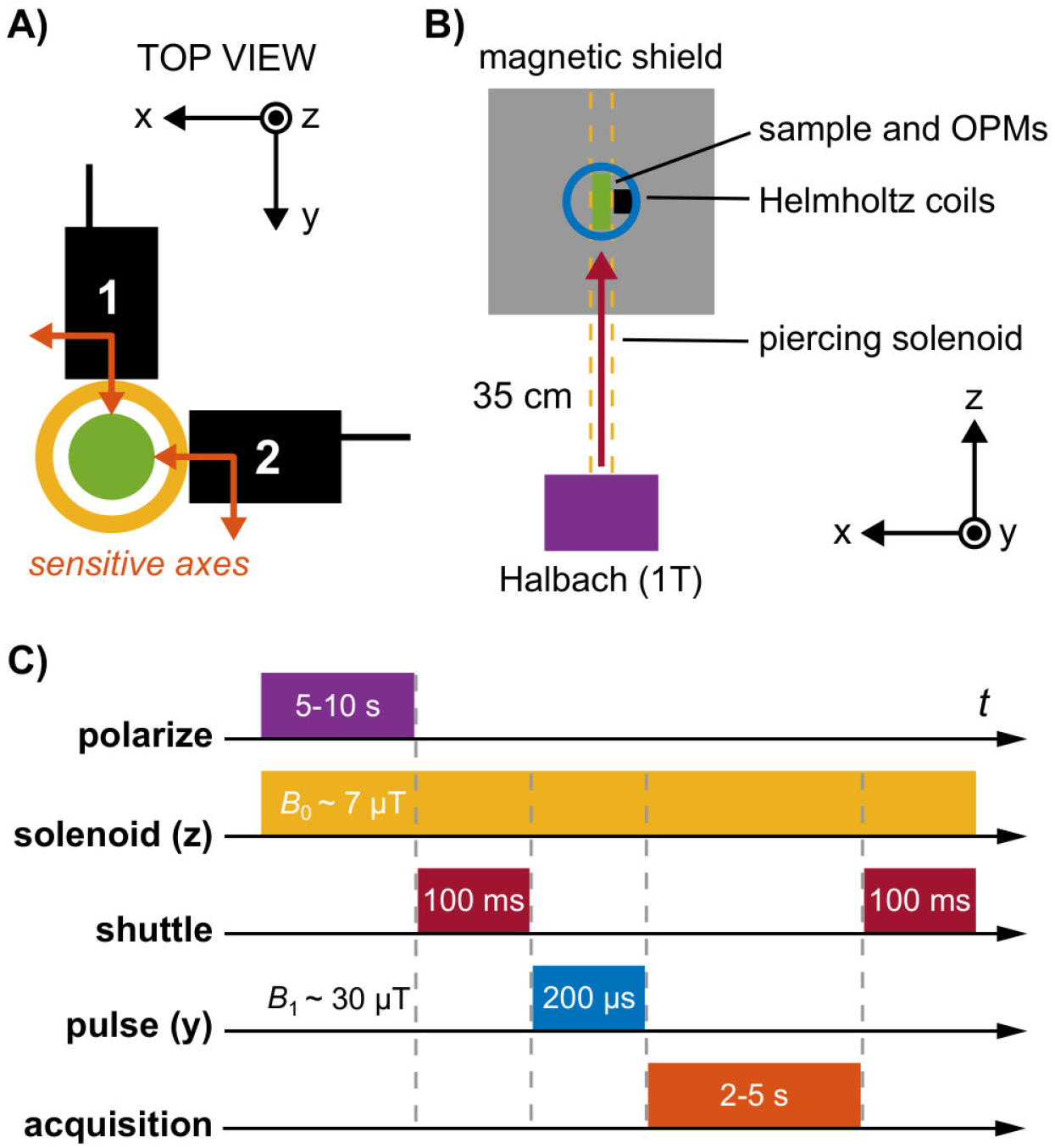
Schematic overview of the relaxometry experiment. **A)** A pair of magnetometers (QuSpin QZFM Gen-2; sensing volume 4 *×* 4 *×* 4 mm^3^, Rb vapor cell), each sensitive along two orthogonal axes indicated with orange arrows, are oriented in the *x*-*y* plane. The center of sensing volume is located 6.5 mm from the tip of the sensor housing (black boxes). Samples to be measured are enclosed in a 2 mL glass vial with outer diameter 11.6 mm (green circle), located in a plexiglass tube around which a solenoid is wound (yellow circle). The outer diameter of the solenoid is approximately 22 mm, so that the minimum offset distance of the sensing volume from the center of the sample is 17.5 mm. **B)** Nuclear spins are first thermally polarized in a 1 T permanent magnet (Halbach array) before being mechanically shuttled into a magnetically shielded environment. There, three orthogonal pairs of Helmholtz coils enable manipulation of the spin states via controlled application of magnetic-field pulses. The piercing solenoid is used both for guiding during shuttling and for generation of a tunable precession field inside the magnetic shield. **C)** The typical experimental protocol uses a guiding magnetic field of 7 μT inside the solenoid (proton precession frequency *∼* 285 Hz). Polarization in the 1 T magnet lasted 5 s for leaf samples and 10 s for water calibration samples, followed by shuttling into the center of the magnetic shield within 100 ms. A 30 μT magnetic-field pulse was then immediately applied in order to rotate *z*-magnetization of the sample into the *x*-*y* plane (*π/*2 pulse), where subsequent free induction decay (FID) of the magnetization signal in the precession field was recorded by the magnetometers. Signal acquisition time was set to 2 s for leaf samples and at least 3 s for water calibration samples.

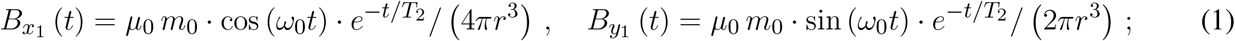

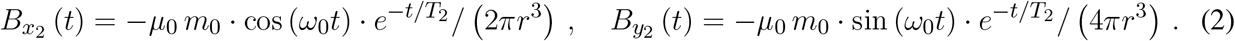

Here, *r* is the offset distance of the center of the sensing volumes from the magnetic dipole (the sample is modeled as a uniformly magnetized sphere); *T*_2_ is the characteristic exponential decay time of the precession signal; *μ*_0_ is the vacuum permeability constant. The initial magnetic-moment amplitude *m*_0_ equals the sample volume times the magnetization of the sample. Note that each measured magnetic-field component may be positive or negative, since vector rather than scalar magnetometers are used. For a spherical 1 mL sample of water polarized at 1 T and room temperature, one can estimate magnetic-field values (at the beginning of the measurement assuming no relaxation losses) on the order of 100 pT, given the experimental offset distance of 17.5 mm. This agrees with experimental data (Figure 2); a complete calculation is provided in Supplementary Material. We note that, fundamentally, Equations (1)–(2) should contain 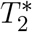 rather than *T*_2_, as we do not employ CPMG or other dynamic decoupling sequences to suppress effects of magnetic-field inhomogeneities at the position of the sample during measurement. However, because the studied leaf samples have intrinsically high relaxation rates (*T*_2_ of 150–300 ms, see Section 3) which are larger than contributions due to inhomogeneity of the solenoid (Figure S8), such simplification is warranted in our case.

**Figure 2.**
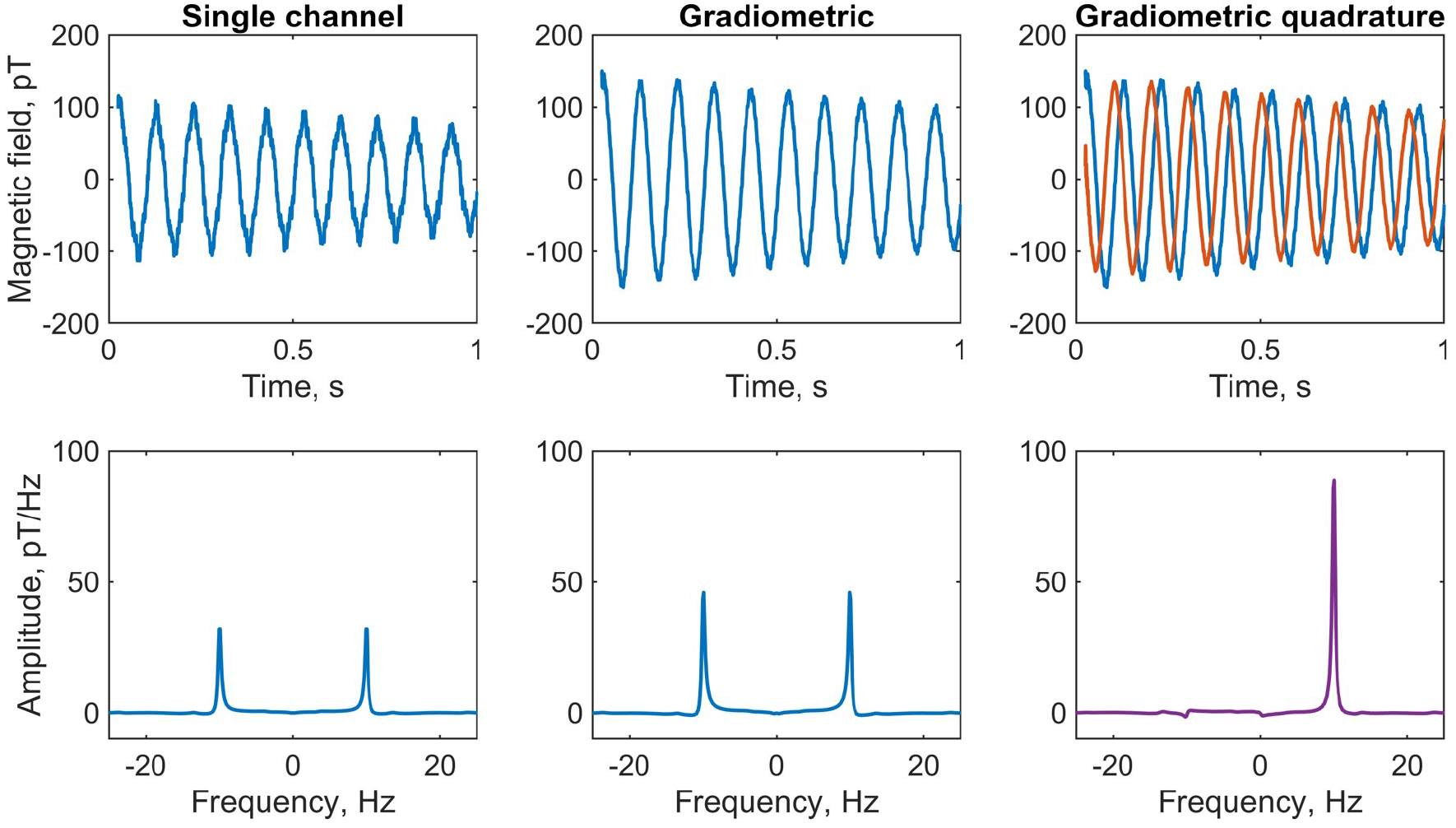
Comparison of water-proton NMR calibration spectra using three different detection modalities. A 2 mL vial of local tap water was measured using the detection geometry and relaxometry protocol depicted in Figure 1, at a 230 nT (10 Hz) precession field. Plots show the average of four scans. Top row, left to right: first second of the free-induction-decay (FID) signal recorded by the *y*-channel of sensor 1; gradiometric FID signal (difference of signals from the *y*-channels of sensors 1 and 2), showing signal enhancement and noise suppression; overlaid *x* (red) and *y* (blue) gradiometric FID signals, which are summed in quadrature as described in the text. Bottom row: corresponding frequency spectra obtained by fast Fourier transform (FFT); the signal at +10 Hz is enhanced 2.9 times in the phased quadrature spectrum (rightmost panel) as compared to a phased single *y*-channel spectrum, and 3.5 times compared to a phased single *x*-channel spectrum. Only a small residual remains at *−* 10 Hz due to imperfections in the quadrature geometry, after correcting for differences in gain of the gradiometric channels.

We see from Equations (1)–(2) and the geometry in Figure 1A that by subtracting the measured fields along the 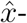 and ŷ-axes, respectively, the signals along each axis add—leading to a signal enhancement of 1.5, assuming identical sensor response—while common-mode noise is canceled. Furthermore, the two gradiometric signals have a relative phase of *π/*2—i.e., are in-quadrature—which becomes useful for signal processing in the frequency domain via Fourier transform. This is the basis of what we have termed the “gradiometric quadrature” detection scheme, used in the work reported here to achieve enhanced (by a factor of *∼*3 compared to single-channel measurement) proton signals in *ex vivo* tree leaves.

The experimental setup, contained in a portable instrument rack, is depicted schematically in Figure 1B. Thermal polarization of nuclear spins is created using a 1 T Halbach magnet with 15 mm bore, which defines the maximum possible diameter of measured samples. After adequate polarization-buildup time in the magnet (5–10 s), rapid (*∼*100 ms) mechanical shuttling of the sample into the magnetic shield (Twinleaf MS-1LF) is performed using an Arduino-controlled stepper motor driving a plastic gear rack, to which a nonmagnetic sample holder is attached. Shuttling occurs inside a double-layer piercing solenoid wrapped around a plexiglass tube and reaching from the top of the Halbach magnet through the magnetic shield. At the center of the shield, a 3D-printed frame of ABS plastic contains three pairs of Helmholtz coils (radius 33 mm) for creation of magnetic-field pulses along the *x*-, *y*-, or *z*-axes to manipulate the nuclear-spin state, as well as two atomic magnetometers (QuSpin QZFM Gen-2) for detection of nuclear-spin signals (Figure 1A). These zero-field sensors can operate in ambient magnetic fields of up to tens of nT, which is readily achieved with the magnetic shield, with the aid of built-in compensation coils. We introduced the piercing solenoid specifically to be able to separate the background field on the sensors from the precession field on the sample. A field on the order of 10 μT may be generated inside the piercing solenoid without compromising sensor operation, as field leakage outside the solenoid is usually below 1%. The range of achievable precession frequencies is ultimately limited by sensor bandwidth—below 500 Hz for the atomic magnetometers used in this work.

Figure 1B shows the typical relaxometry protocol for the experiments reported here. After spin polarization and shuttling, immediate application of a *π/*2-pulse in the *y*-direction rotates the bulk *z*-magnetization into the *x*-axis, where it subsequently precesses about the leading *z*-field of amplitude *B*_0_ at an angular frequency *ω*_0_ given by the proton gyromagnetic ratio 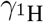 according to 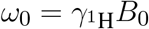, where 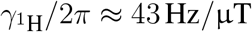. This Larmor precession gives rise to a free-induction-decay (FID) signal which is acquired by the magnetometers. For proton spins (positive sign of the gyromagnetic ratio), precession is “left-handed”, occurring clockwise about the applied magnetic field [34].

Experimental timing and control as well as detector readout were implemented in Labview using NI TTL-pulse and data-acquisition cards. To maximize the SNR of the measured FID signal, a number of experimental parameters were iteratively optimized. These include: polarization time, solenoid field (proton precession frequency), shuttling time and speed/acceleration, pulse duration and amplitude, and acquisition time. Various calibration data, along with photos and further details of the apparatus, may be found in Supplementary Material.

### 2.2 Data processing

During acquisition of an FID according to the protocol in Figure 1B, the analog voltage outputs of all four magnetometer channels are recorded at a user-defined sampling rate, usually 2 kHz, for subsequent analysis. Although the sensors are always calibrated (i.e., ambient magnetic fields internally compensated) with the sample in the measurement position prior to each experiment, application of the magnetic-field pulse drives the sensors out of their sensitive range for some tens of milliseconds. Thus, initial data points must be discarded in post-processing, resulting in an effective linear phase shift of the recorded oscillating signal. Because each magnetometer channel provides a vector measurement—sensitive to the sign of the magnetic field, in contrast to a scalar sensor—the handedness of spin precession may be deduced from a single-channel time trace, if it is possible to reconstruct the true phase of the FID. However, the quadrature detection scheme allows us to determine the handedness without having to reconstruct the phase.

As an illustration of the analysis procedure, Figure 2 shows calibration data from a liquid water sample at a proton precession frequency of 10 Hz, obtained by supplying a current of 54 μA to the piercing solenoid. The first 25 ms of data, corresponding to the first 50 points of the FID sampled at 2 kHz, have been discarded to remove post-pulse artifacts which would otherwise adversely affect the spectral lineshape and baseline. Plots show the average of multiple (here, four) scans, where a linear trend has been removed from each individual raw time trace prior to averaging. Initially, we convert from voltage to magnetic-field units to obtain four time series associated with the four magnetometer channels, which we denote *x*_1_, *y*_1_, *x*_2_, and *y*_2_, following Equations (1)–(2). The two gradiometric channels are subsequently constructed as *x* = *x*_1_ *− x*_2_ and *y* = *y*_1_ *− y*_2_. By examining the single-channel and gradiometric time traces in the context of the detection geometry (Figure 1A), we can confirm that the spin signal was initialized along 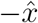 and is precessing clockwise in the *x*-*y* plane, consistent with a precession field along 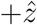. Furthermore, reduction of noise via gradiometry is clearly visible.

The gradiometric-quadrature channel can be constructed from the two gradiometric channels as *x*+*iy* (see Supplementary Material for more details). In our experimental geometry, the time-dependent quadrature signal may be thought of as a vector rotating clockwise in the complex plane defined by a real *x*-axis and imaginary *y*-axis. Assuming perfect quadrature geometry, when a standard Fourier transform is performed to convert the signal into the frequency domain, the resulting spectrum should contain a resonance only at +10 Hz and not at *−*10 Hz.

In the time domain, the processed complex gradiometric-quadrature signal is written as

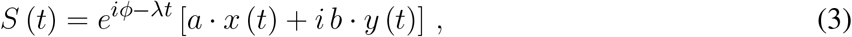

where *x* (*t*) and *y* (*t*) are the gradiometric signals, *ϕ* is an overall phase selected such that the real part of the frequency spectrum has an absorptive peak (see below), and *λ* is an optional apodization factor [35]. The numerical coefficients *a* and *b* are defined so that *a* + *b* = 2, and may be adjusted to account for possible differences in gain between the two channels—usually only a few percent, based on suppression of the residual negative-frequency peak. Prior to the Fourier transform, zeros may be added to the end of the time series defined by Equation (3) (zero-filling) in order to increase the spectral resolution.

All frequency spectra throughout this manuscript are plotted such that the *y*-axis has units of pT/Hz or fT/Hz, depending on the signal strength. These units arise from the discrete Fourier transform (DFT) used to convert data from the time domain to the frequency domain and subsequent data processing, as explained in detail in Supplementary Material.

The exact SNR enhancements attainable by the gradiometric quadrature method depend strongly on performance of the individual sensors, which may vary between experiments, as well as non-common-mode systematic noise. In our experience with water calibration samples, compared to single-channel spectra, SNR was enhanced *∼* 30–50% via gradiometry alone and *∼* 40–60% via the gradiometric quadrature method. This indicates not only that gradiometry is effective in terms of noise suppression, but also that the quadrature approach is more beneficial than simply summing the positive-frequency and negative-frequency peaks in a traditional “mirrored” non-quadrature spectrum. See Supplementary Material for quadrature simulations, gradiometer sensitivity data, and further details about phasing of quadrature signals. We expect that gradiometric quadrature detection can be especially beneficial for situations in which sensitive magnetometers operate in unshielded environments—if the contribution from common-mode noise dominates the contribution from uncorrelated noise at the positions of two sensors, total measurement noise will be significantly suppressed.

### 2.3 Leaf harvest and sample preparation

A description of all 19 tree-leaf samples included in our study is provided in Table 1. For identification purposes, samples from each genus were numbered in order of preparation/measurement date. We note that, due to seasonal availability, evergreen (spruce, pine, and yew) leaves were collected from late February to early April, while deciduous (hazel, cherry, beech, and oak) leaves were collected from early April to late May. Sample collection occurred as close as possible to the time of first measurement. All samples were sourced from the Eurasian arboretum of the Mainz Botanical Garden, which informed the specific choice of species and/or cultivar, although we purposely selected a diverse range of common genera.

**Table 1.**
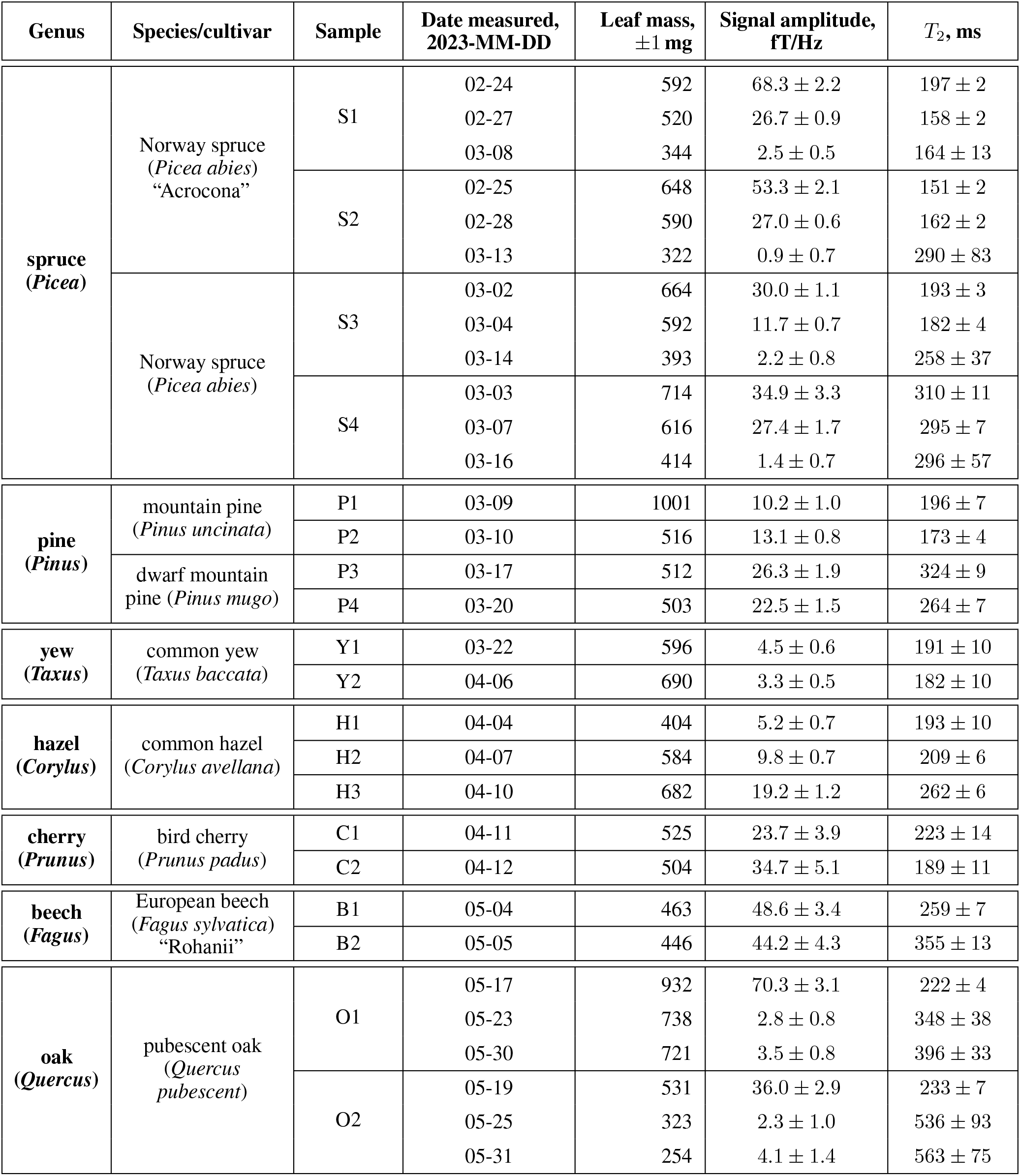
Overview of the leaf measurement campaign, in which samples from seven different tree genera were studied. Branchlets were freshly harvested in the Mainz Botanical Garden and transported to the lab in a beaker of distilled water for immediate sample preparation and measurement. Intact leaves were carefully dried and packed into a 2 mL closed glass vial, which was weighed and inserted into the experimental apparatus for a 13 h measurement (4096 scans). For dehydration studies (spruce and oak), sample vials were stored uncapped between measurements in a plant growth chamber at 22^*°*^C.

During harvest, a branchlet containing sufficient leaf coverage was cut from the branch tip of the tree donor and immediately placed in a glass beaker partially filled with distilled water such that the cut end of the branchlet was submerged, as shown in the photo inset of Figure 3. Sample preparation in the laboratory proceeded as follows. Leaves or needles were gently removed from the branchlet, dried with a clean tissue to remove any excess moisture, and packed into a pristine glass shell vial (BGB SV2ML) with outer diameter 11.6 mm and interior volume 3.3 cm^3^. Spruce, pine, and yew needles were placed lengthwise vertically into the vial, whereas the hazel, cherry, beech, and oak leaves had to be rolled or folded. Care was taken to avoid tearing or otherwise damaging leaves during sample preparation, thereby preserving the original structure of the plant tissue, while fitting as many leaves or needles as possible into the tubular vial (Figure 3 photo inset). Due to leaf geometry, it was not possible to achieve completely uniform density of leaf material in the vial, particularly for larger deciduous leaves. Each individual vial and its plastic cap were weighed with a digital scale before and after leaf insertion to extract the total leaf mass. Vials were capped from the moment of leaf insertion and throughout each 13 h experiment in the relaxometry setup, to prevent dehydration of the sample. After each experiment, the capped vial was removed from the setup, uncapped to release any trapped water vapor, recapped and weighed again. We found that post-measurement mass was always within a few mg of pre-measurement mass (Table 1), suggesting that leaf dehydration during the experiment was negligible.

**Figure 3.**
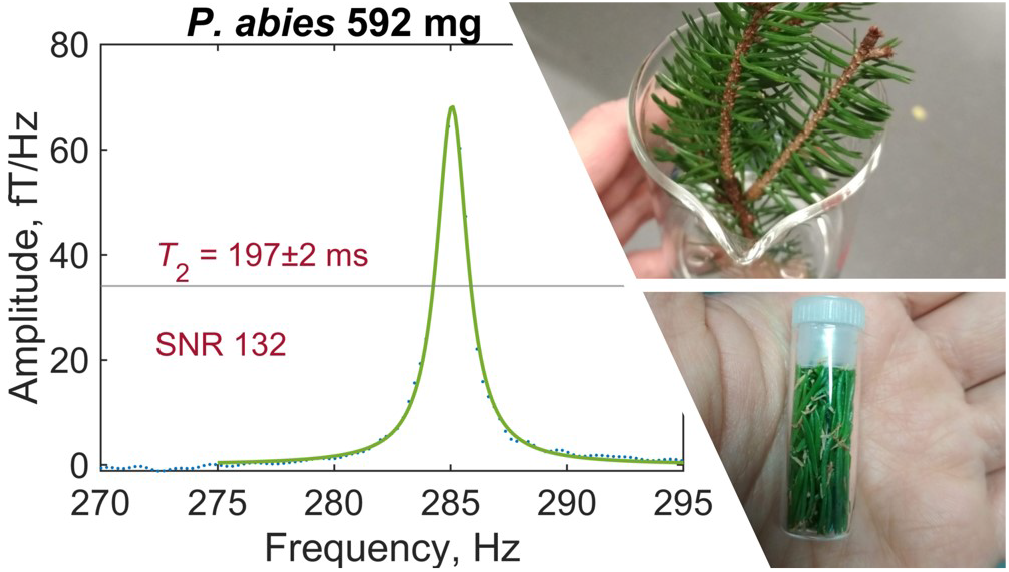
Example of a typical leaf spectrum recorded by gradiometric quadrature detection (spruce sample S1). Water-proton signal has been fitted with a Lorentzian and phased such that fit residuals are minimized—see text for details of data analysis. Inset photos show a freshly harvested spruce branchlet (top) and a prepared sample of spruce needles (bottom).

For the dehydration investigation, one evergreen genus (spruce) and one deciduous genus (oak) were selected; each sample was measured three different times, on the dates indicated in Table 1. Between experiments, the sample vial was stored uncapped in an on-site plant growth chamber (poly klima PK-520, 22^*°*^C, 12/12 h light/dark cycle, no humidity control). Because the vial was open only at one end, uniform dehydration of the leaf content could not be ensured. However, visible inspection of leaf color as well as the usual weighing procedure indicated that water loss had occurred during each storage period. It was critical to ensure that no condensation collected inside the vials at any stage of dehydration or measurement preparation, as this could introduce an additional spurious water-proton signal not originating from water protons contained in the leaves themselves.

### 2.4 Leaf measurements

Each leaf measurement was conducted under identical experimental conditions to produce a proton-NMR spectrum as in Figure 3. A resonance frequency of around 285 Hz, corresponding to a 7 μT precession field with 1.5 mA applied to the piercing solenoid, was selected to avoid lower-frequency noise while remaining with the sensitive bandwidth of the magnetometers (see Supplementary Material). In the plotted spectrum, linear background was removed in the region 260–320 Hz and a Lorentzian fit (three-parameter Lorentzian function with constant term) was performed using the *lorentzfit* script in Matlab. As expected, the experimental lineshape is well-fitted by a Lorentzian, as it results from an exponentially decaying signal. Phasing of the quadrature signal was optimized by minimizing the root mean square error (RMSE) of the fit. Using the calculated fit parameters and errors thereon, the fit amplitude and linewidth (FWHM, full width at half maximum) were extracted. Typical fit amplitudes were on order 1 to 10 fT/Hz—at least three orders of magnitude smaller than for pure water samples of similar volume—after averaging over 4096 scans (repetitions of the measurement protocol). The relaxation time *T*_2_ is related to the FWHM Δ as 1*/*(*π*Δ). Reported SNR values were obtained from the ratio of fit amplitude to the standard deviation of spectral noise in the region 265–275 Hz. To avoid artificial broadening of the spectral line, no apodization was used, i.e. *λ* = 0 in Equation (3); instead, a 300 Hz low-pass filter was applied to each spectrum for lineshape correction of the averaged signal. Measured proton-signal amplitudes and *T*_2_ times are recorded for all 35 leaf experiments in Table 1. Complete fitted spectra, along with example analysis code and further details of the analysis protocol, may be found in Supplementary Material.

In principle, multiple mechanisms may affect the spectral linewidth. These include magnetic-field gradients at the location of the sample, combined effects of different relaxation mechanisms, and possible contributions from non-water protons. Due to the significant differences in linewidth between liquid and leaf samples, we conclude that field inhomogeneity at the location of the sample is negligible, and therefore we report *T*_2_ rather than 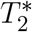 times in this work. Because experimental parameters remained unchanged over the course of the leaf study and all data underwent an identical analysis procedure, direct comparison of signal amplitudes and *T*_2_ times is possible. Comparing the leaf results to typical water calibration data, we find that *T*_2_ times in fresh leaf samples are an order of magnitude shorter than in liquid water samples of similar volume.

## 3 RESULTS

Figures 4 and 5 provide graphical representations of the results reported in Table 1. In Figure 4A, we see that the normalized signal amplitude of the spectra obtained from freshly prepared leaf samples varies significantly by tree genus. This may indicate different water-storage capacities of leaves from different genera. It is interesting to note that packing more leaf matter into the sample vial to increase the overall sample mass did not necessarily increase the normalized signal amplitude, for leaves of the same species. This could be attributed to two effects: (1) non-uniformity of the packing of leaf matter inside vials and (2) demagnetization effects due to the fact that samples are cylindrical rather than spherical. Due to leaf shape, distribution of leaf material in the sample vial is not necessarily uniform and this is generally worse for deciduous trees than for evergreen trees, due to the needle-like form of the latter. We observe that evergreen tree leaves fill up vials more uniformly. Demagnetization field effects due to cylindrical sample geometry can result in lowered signal for some samples compared to others (Table 1). Future studies are warranted to investigate the effects of sample geometry and uniformity of leaf matter on the magnitude of observable NMR signals and the precise amount of water giving rise to them. In contrast to the amplitude results, we see from Figure 4B that average measured *T*_2_ times do not appear to vary significantly among the studied tree genera.

**Figure 4.**
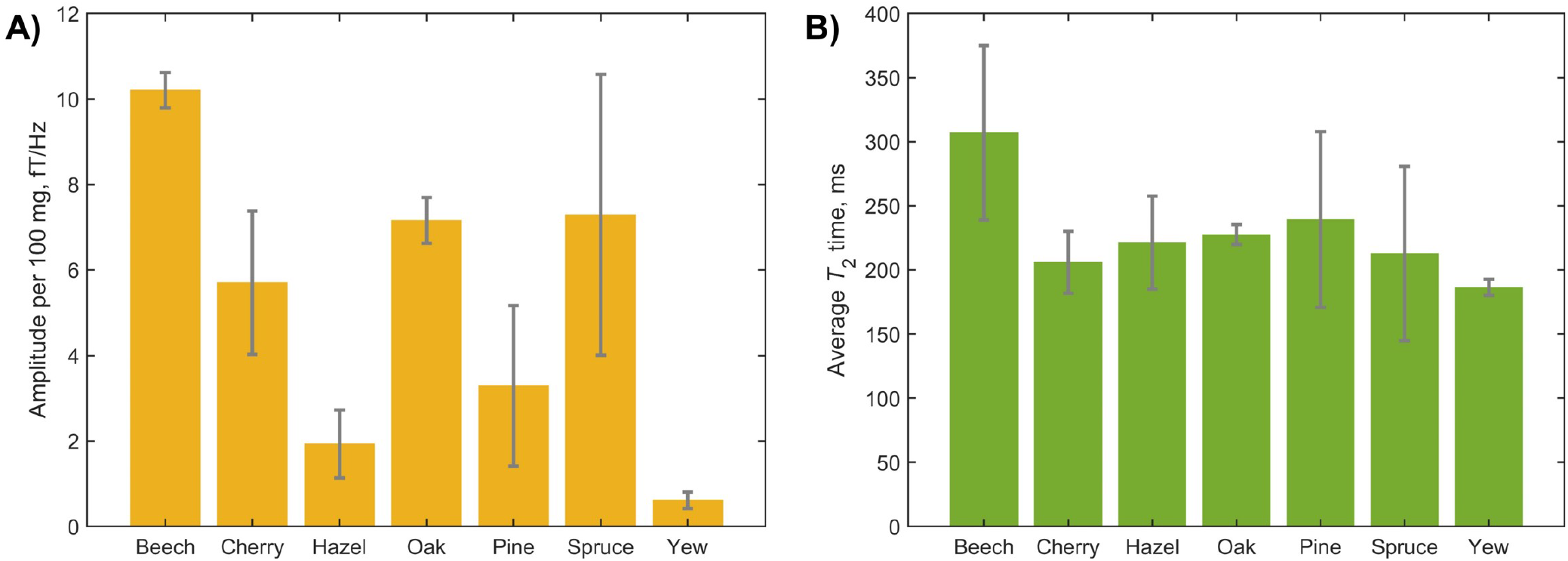
Results of the cross-species leaf-relaxometry study encompassing 19 fresh leaf samples. **A)** Average normalized measured water content from the different tree genera. Signal amplitudes were extracted from Lorentzian fits of the measured spectra; error bars indicate the standard deviation of signal amplitude for each genus. Note that the spruce and pine data encompass multiple species or cultivars (Table 1). **B)** Average *T*_2_ times, extracted from Lorentzian fits of the measured spectra; error bars indicate the standard deviation for each genus.

**Figure 5.**
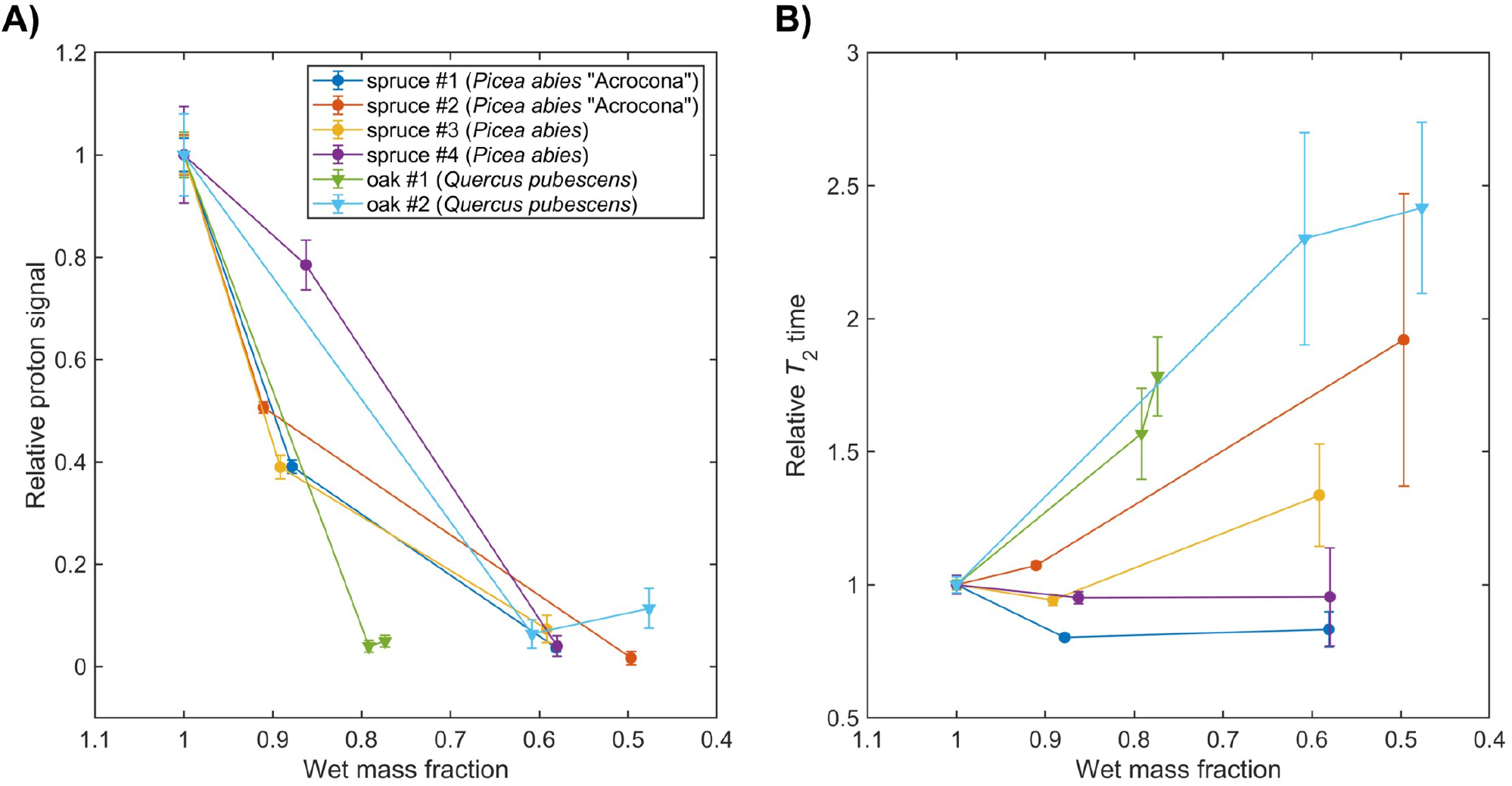
**A)** Tracking water content with dehydration in spruce (*P. abies*) and oak (*Q. pubescens*) leaves, see Table 1. Signal amplitudes and errors were extracted from Lorentzian fit parameters of the measured spectra; lines connecting data points are a guide to the eye. **B)** Tracking proton *T*_2_ relaxation times with dehydration. Relaxation times and errors were extracted from Lorentzian fit parameters of the measured spectra; lines connecting data points are a guide to the eye.

In Figure 5, showing the results of the spruce/oak dehydration study, two trends can be observed. First, we observe an overall decrease in signal amplitude as sample mass decreases due to water loss, indicating that our experiments are primarily sensitive to water protons, as expected. However, we note that the dependence on sample mass is not entirely linear—this may be related to inhomogeneous dehydration of the sample, as mentioned above, as well as possible signal contributions from non-water protons. Second, an overall increase of *T*_2_ times, i.e. narrowing of the water-proton peak, is observed as sample mass decreases. This trend is more pronounced for the oak samples, even accounting for increased uncertainty on *T*_2_ due to reduction of SNR with dehydration. Notably, *T*_2_ increases most dramatically for oak samples, as compared to spruce.

## 4 DISCUSSION

In this study, we showcased several key findings, including the noninvasive and nondestructive measurement of water signals in intact *ex vivo* plant parts using a proton relaxometry protocol at hypogeomagnetic field. Additionally, we achieved signal-to-noise ratio enhancement of weak biological NMR signals from non-solution samples by employing a gradiometric quadrature detection scheme, especially useful in a future deployment of this technology in the field. Our research involved a comparative investigation of water-proton signals and *T*_2_ relaxation in 19 tree-leaf samples, encompassing samples from seven genera, eight species, and nine cultivars. With this, we demonstrated sensitivity to the evolution of water-proton signals and *T*_2_ relaxation times through repeated measurements of dehydrating leaf samples.

The experiments reported here were intended as a proof-of-principle of the above, and have not yet attempted to answer specific biological questions. Nonetheless, the preliminary results displayed in Figures 4–5 already contain information which suggests future lines of relaxometry research with tree leaves. For example, the observed differences in normalized water-proton signal amplitude among different genera and species/cultivars may motivate further large-sample-size studies of water-storage capacity and possible seasonal variations. By contrast, the relative uniformity of measured *T*_2_ times in all fresh leaf samples indicates that, at least in the hypogeomagnetic field regime, water-proton relaxation in leaf tissue is dominated by mechanisms common to the studied tree types. The observed tendency toward lengthening of *T*_2_ times (narrowing of the proton precession peak) with leaf dehydration, particularly in the measured oak samples, may seem contradictory to intuition—if one expects dehydration and tissue death to further constrain molecular motion and lead to broadening of the spectral feature. However, our result appears to be consistent with previous benchtop relaxometry studies where leaf senescence was correlated with changes in water distribution at the cellular level as well as lengthening of *T*_2_ components [13, 14]. Thus, we hope that our tree-leaf dehydration study will help open to the door to further relaxometry-enabled research on drought stress and tolerance in the context of forestry and agriculture.

Further improvements to the experimental setup will enable the affordable atomic-magnetometer based relaxometry device to achieve the functionality of commercial benchtop spectrometers for biological applications. These refinements may include implementation of spin-echo pulse sequences, SNR enhancements via suppression of low-frequency noise and optimization of the shuttling field profile, and shimming (field compensation) of stray magnetic fields and gradients. Relaxation-dispersion studies (measuring relaxation times as a function of field) may also reveal further information about water-storing structures [36, 37, 38]. Instrumentation such as custom magnetometers tailored to plant samples—with reduced standoff distance and surface temperature—will improve biocompatibility, and the use of Earth-field magnetometers would even enable unshielded measurements (see [39] and references therein). The shielded regime is itself of fundamental interest, for example in studying properties of biological tissues under hypogeomagnetic conditions such as those encountered during long-distance spaceflight. Relaxometry studies of systems in which NMR signals originate from molecules other than water are also valuable, since other relaxation mechanisms can be involved [40]. While future experiments need not be limited to relaxometry of protons only, NMR-enabled investigation of plant water dynamics is highly warranted, particularly in ultralow and hypogeomagnetic regimes.

## AUTHOR CONTRIBUTIONS

Conceptualization: D.A.B. and A.M.F. Data curation and analysis: A.M.F. Funding acquisition: D.A.B. Investigation and methodology: A.M.F., P.P., and D.A.B. Resources: D.A.B. Software: P.P. and A.M.F. Supervision: D.A.B. Visualization and writing—original draft: A.M.F. Writing—review & editing: A.M.F., P.P., and D.A.B.

## ACKNOWLEDGMENTS

We thank Prof. Dmitry Budker and Erik Van Dyke for stimulating discussions and feedback, and acknowledge contributions from Kirill F. Sheberstov, Liubov Chuchkova, Oleg Tretiak, and Raphael Kircher in initial development of the experimental setup. The Halbach magnet used for spin polarization was designed by Dr. Peter Blümler. We thank the Mainz Botanical Garden Arboretum for providing samples for this study.

## DATA AVAILABILITY STATEMENT

All datasets generated for this study are available upon request.

