## Supplemental Material for "Proton Relaxometry of Tree Leaves at Hypogeomagnetic Fields"

1 Photographs and apparatus 1
2 Expected magnetic-field amplitudes 1
3 Gradiometric quadrature detection: Simulations and signal processing 3
4 Sensor performance 3
5 Calibration of experimental parameters 3
6 Leaf data 3
7 Data analysis 3

### 1 PHOTOGRAPHS AND APPARATUS

Figure S1 shows a photo of the experimental setup as well as a selection of measured samples. The following hardware is included in the setup:

- *Experimental timing and data acquisition*—NI cDAQ-9189 ethernet chassis, two NI 9401 TTL-pulse cards with NI 9924 adapters, NI 9215 BNC acquisition card;
- *Spin polarization*—custom 1 T Halbach magnet produced in-house;
- *Mechanical shuttling*—Arduino Uno, Sanyo Denki StepSyn stepping motor 103H7123-5640 with driver, RS PRO plastic gear rack and aluminum rail;
- *Magnetic shielding*—Twinleaf MS-1LF ferrite magnetic shield with CS121 current supply;
- *Spin manipulation*—homemade copper double-layer piercing solenoid (inner diameter 20 mm, height 50 cm, resistance $11.5\,\Omega$, calibration 4.5 nT/μA) wound around an acrylic tube (Acrylhaus) and connected to DM Technologies Multichannel Current Source, homemade 3D-printed ABS-plastic coil frame holding three orthogonal Helmholtz coil pairs (radius 33 mm, resistance $0.9\,\Omega$, 10 copper windings, calibration 256 nT/mA) connected to Mini-Circuits switch ZASWA-2-50DRA+ and three Basetech BT-305 DC power supplies;
- *Detection*—two QuSpin Zero-Field Magnetometers (QZFM Gen-2) also contained in the coil frame.

Many of the commercial electronic items could be replaced with lower-cost or homemade alternatives; our choice of equipment was largely informed by what was already available in our labs.

### 2 EXPECTED MAGNETIC-FIELD AMPLITUDES

For a pure water sample polarized at $B_\mathrm{p} = 1$ T and room temperature $T = 293\,\mathrm{K}$, the expected thermal polarization of protons is [1, 2]

$$P_\text{therm} = \frac{\gamma_{^1\mathrm{H}}\,\hbar\,(I+1)\,B_\mathrm{p}}{3\,k_\mathrm{B}T} = \frac{\gamma_{^1\mathrm{H}}\,\hbar\,B_\mathrm{p}}{2\,k_\mathrm{B}T}\,, \tag{S1}$$

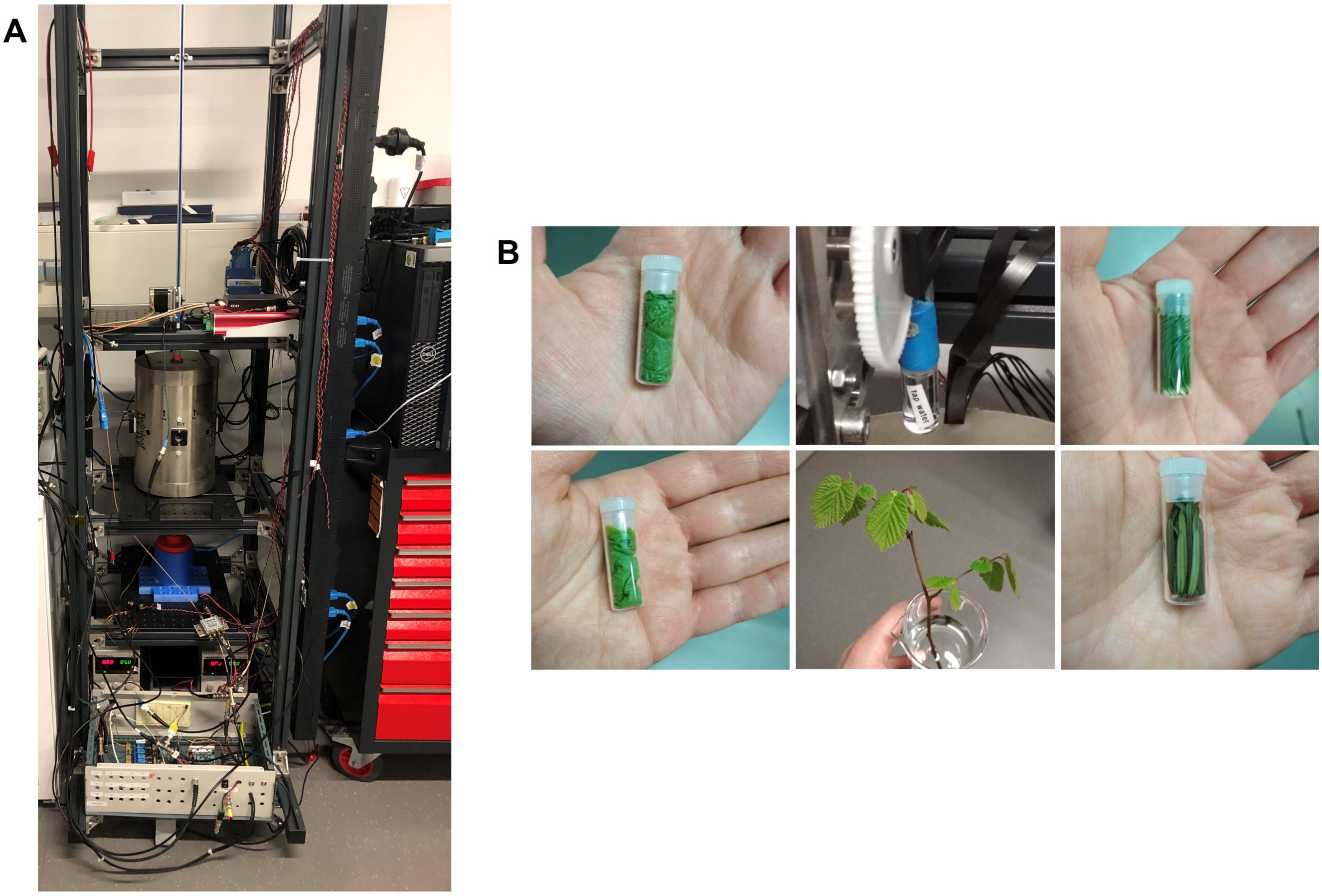

**Figure S1.** Photo collage of the relaxometry setup and samples. **(A)** Experimental apparatus, cf. Figure 2a in the main text. **(B)** Clockwise from top center: water calibration sample, pine sample, yew sample, harvested hazel branchlet, cherry sample, hazel sample.

where $\gamma_{^1\mathrm{H}}/2\pi = 42.6$ MHz/T and $I = 1/2$. This calculation yields

$$P_{\text{therm}} = \frac{\left(42.6 \times 10^{6}\,\mathrm{Hz}\cdot\mathrm{T}^{-1}\right)\left(6.63 \times 10^{-34}\,\mathrm{J}\cdot\mathrm{s}\right)(1\,\mathrm{T})}{2\left(1.38 \times 10^{-23}\,\mathrm{J}\cdot\mathrm{K}^{-1}\right)(293\,\mathrm{K})} = 3.5 \times 10^{-6}\,. \tag{S2}$$

Given the molar concentration of water protons, $C = 110$ M, the amplitude of sample magnetization (magnetic moment per unit volume) can then be estimated as

$$\begin{aligned} M &= C \cdot \gamma_{^1\mathrm{H}}\hbar I \cdot P_{\text{therm}} \qquad \text{(S3)} \\ &= \left(110\,\mathrm{mol}\cdot\mathrm{L}^{-1}\right)\left(6.02 \times 10^{23}\,\mathrm{mol}^{-1}\right)\left(42.6 \times 10^{6}\,\mathrm{Hz}\cdot\mathrm{T}^{-1}\right)\left(6.63 \times 10^{-34}\,\mathrm{J}\cdot\mathrm{s}\right)\left(\frac{1}{2}\right)\left(3.5 \times 10^{-6}\right) \\ &= 3.3 \times 10^{-6}\ \mathrm{A}\cdot\mathrm{m}^{2}\cdot\mathrm{L}^{-1} = 3.3 \times 10^{-6}\ \mathrm{A}\cdot\mathrm{m}^{-1}\,. \end{aligned}$$

For a sample of volume $V = 1$ mL, the magnetic moment is then $m = \mathrm{M}\cdot\mathrm{V} = 3.3 \times 10^{-9}\ \mathrm{A}\cdot\mathrm{m}^{2}$.

Let us assume that the sample is a uniformly magnetized sphere, giving rise to a pure dipole magnetic field outside. Referring to Equations (1)–(2) in the main text and ignoring any relaxation mechanisms, the maximum magnetic-field amplitude measured at a standoff distance 17.5 mm from the center of the sample is

$$B = \frac{\mu_0\, m}{2\pi r^3} = \frac{\left(1.26 \times 10^{-6}\,\mathrm{N \cdot A^{-2}}\right)\left(3.3 \times 10^{-9}\,\mathrm{A \cdot m^2}\right)}{2\pi\,(0.0175\,\mathrm{m})^3} \approx 120\,\mathrm{pT}\,. \tag{S4}$$

### 3 GRADIOMETRIC QUADRATURE DETECTION: SIMULATIONS AND SIGNAL PROCESSING

Figures S2 and S3 correspond to Figure 3 in the main text—Figure S2 illustrates processing of a simulated time trace, while Figure S3 shows the effect of phasing on experimental results.

### 4 SENSOR PERFORMANCE

Gradiometer sensitivity is presented in Figure S4—both channels exhibit a noise floor of around 100 fT/$\sqrt{\mathrm{Hz}}$ within the bandwidth of the magnetometers. Feedback control of the atomic-vapor-cell heater is turned off during measurement, as it is affected by the DC magnetic-field pulses and may introduce additional low-frequency noise which negatively impacts measured lineshapes.

### 5 CALIBRATION OF EXPERIMENTAL PARAMETERS

Various calibration data are included here, showing optimization of the following: shuttling distance above the polarizing Halbach magnet (Figure S5), duration of the applied DC magnetic-field pulse (Figure S6), polarization time in the magnet (Figure S7), and choice of proton-spin precession frequency (Figure S8).

We expect a $\pi/2$ DC pulse, which exploits Larmor precession to rotate magnetization by 90°, to fulfill the condition

$$2\pi\gamma_{^1\mathrm{H}}\, B_\mathrm{y}\, \tau_\mathrm{p} = \frac{\pi}{2}\,. \tag{S5}$$

Here $B_\mathrm{y}$ is the pulse amplitude, held constant at 30 μT in our setup, and $\tau_\mathrm{p}$ is a variable pulse duration. Given the proton gyromagnetic ratio $\gamma_{^1\mathrm{H}}/2\pi \approx 43\,\mathrm{Hz}/\mu\mathrm{T}$, we expect an optimal pulse duration of around 200 μs, which agrees well with the experimental data plotted in Figure S6.

Best practice to obtain optimized nuclear magnetic resonance (NMR) spectra without lineshape correction involves polarizing spins for $\sim 3$ times the characteristic magnetization decay time $T_1$, and acquiring the free-induction-decay (FID) signal for $\sim 5$ times the $T_2$ time. However, reduction of the experimental duty cycle is desirable in order to more efficiently increase the number measurement averages with associated signal-to-noise ratio (SNR) enhancement. Note that $T_1$ times may depend on magnetic-field strength—see Figure S7.

### 6 LEAF DATA

Figures S9–S12 show fitted water-proton signals from all measured leaf samples, cf. Table 1 in the main text. Oscillations in the spectral baseline are attributed to drifts in magnetometer response over the course of the 13 h data run, which lead to artifacts in the average time traces.

### 7 DATA ANALYSIS

Processing of the experimental data is performed as follows. We begin with a discrete time series $S\,(j)$, a vector containing $z$ data points, where $z$ is a positive integer value usually chosen to be a power of 2. The

DFT is carried out pointwise for each point $j$ using a standard fast Fourier transform (FFT) algorithm in order to produce a new vector $S(k)$:

$$S(k) = \sum_{j=1}^{z} S(j)\, e^{[(-2\pi i)(j-1)(k-1)]/z}\,. \tag{S6}$$

Note that because each point of $S(k)$ is a sum over all points in the time series, the amplitude scales with $z$ and has units of magnetic field. This amplitude may be normalized by rescaling the vector as $S(k)/z$. Although Equation (S6) provides a measure of frequency, it is agnostic with respect to $x$-axis units, and the output vector $S(k)$ has the same length, $z$, as the input vector $S(j)$. In order to plot the spectrum as a function of frequency, we define a frequency axis from $-f_s$ to $f_s$, where $f_s$ is the sampling rate of acquisition, with frequency spacing $f_s/z$ between points. Furthermore, the zero-frequency components of $S(k)$ must be shifted to the center of the spectrum. Example code for carrying out these procedures in Matlab is provided.

According to NMR convention, the integral of a signal in frequency space should equal the initial amplitude of the FID and therefore have units of magnetic field; a convenient normalization sets the integral to 1, such that the mirrored peaks in non-quadrature spectra (Figure S2) each have integral 0.5. This is only possible if the $y$-axis of the frequency spectrum has units of magnetic field per Hz. Such units do indeed arise in the analytical case of a continuous FT, in which an integral rather than a summation is performed over the time-domain signal. In order to approximate the continuous FT using a numerical DFT, we divide the frequency signal $S(k)$ by the frequency spacing or "bin size" $f_s/z$, which yields the desired units and integration properties.

Lineshape fitting of the gradiometric quadrature signal is performed using a three-parameter Lorentzian function

$$L(\nu) = \frac{p_1}{(\nu - p_2)^2 + p_3} + c\,, \tag{S7}$$

where $\nu$ is frequency, $p_1$, $p_2$, and $p_3$ are the fit parameters, and $c$ is a constant. Spectral properties may be extracted from the fit parameters as follows: $p_2$ is the central peak position in Hz, $p_1/p_3$ is the fit amplitude, and $T_2 = 1/(2\pi\sqrt{p_3})$—equivalent to $1/(\pi\Delta)$, where $\Delta$ is the full width at half maximum (FWHM). In order to optimize phasing of the spectrum as described in the main text, the phase $\phi$ is varied in steps of $\phi/20$ until the root mean square error (RMSE) calculated from fit residuals is minimized.

Further details of analysis steps may be found in the provided annotated Matlab analysis notebook 'AnalyzeLeaf.m', which was used to produce the leaf spectra plotted in Figures S9–S12.

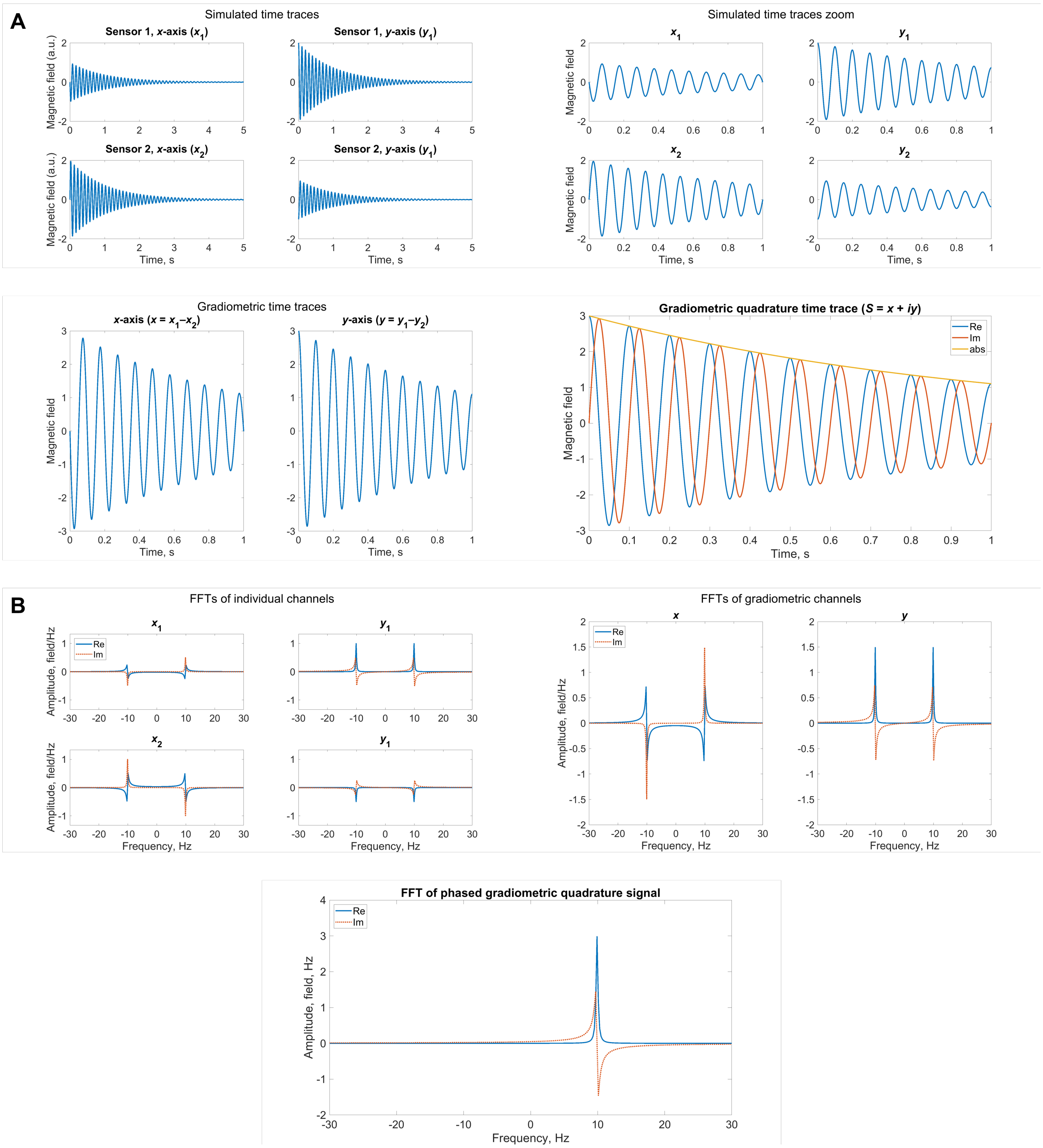


**Figure S2.** Simulations of magnetic-field measurements of a magnetic dipole precessing at 10 Hz in the $x$-$y$ plane, following the detection geometry in Figure 1 of the main text. Here we assume that magnetization has been initialized along $\hat{y}$ prior to measurement and relaxes with a $T_2$ time of 1 s. **(A)** Normalized time traces showing the free-induction-decay (FID) signal as seen by the four individual magnetometer channels (top row), the two gradiometer channels (bottom left), and the gradiometric quadrature channel (bottom right). **(B)** Corresponding frequency spectra obtained by fast Fourier transform (FFT). An overall phase of $-\pi/2$ has been applied to the gradiometric quadrature time trace such that the real and imaginary parts of the resulting FFT (bottom panel) are absorptive and dispersive, respectively.

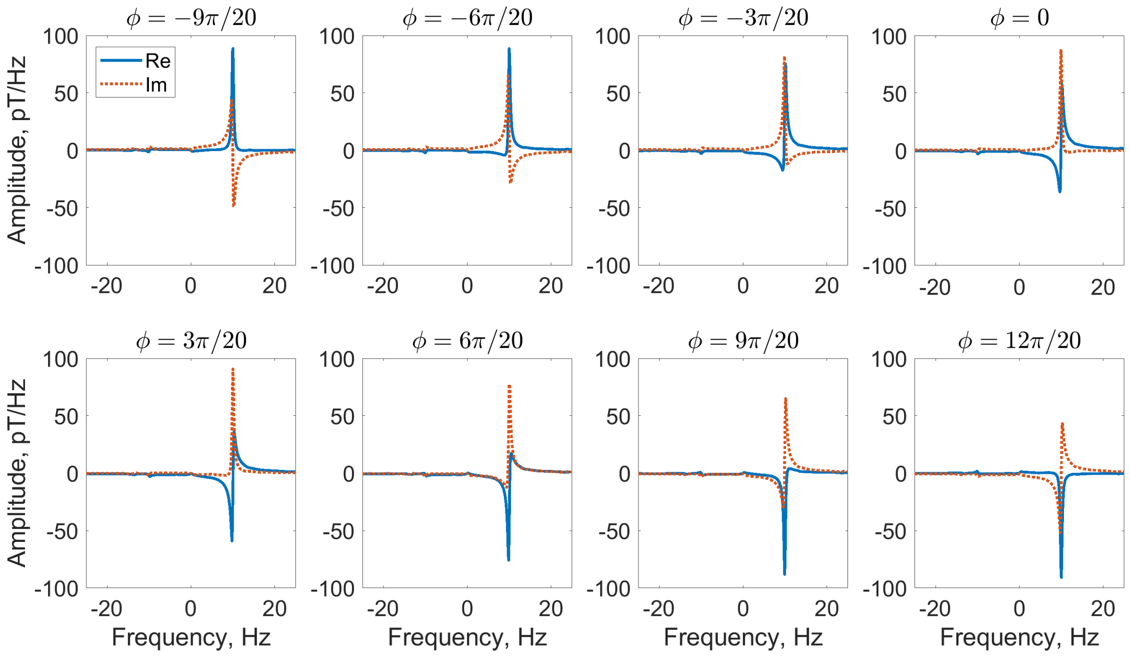


**Figure S3.** Phase sensitivity of the water calibration spectrum, cf. Figure 3 in the main text. To produce these plots, the measured gradiometric quadrature signal was multiplied by a variable overall phase $e^{i\phi}$.

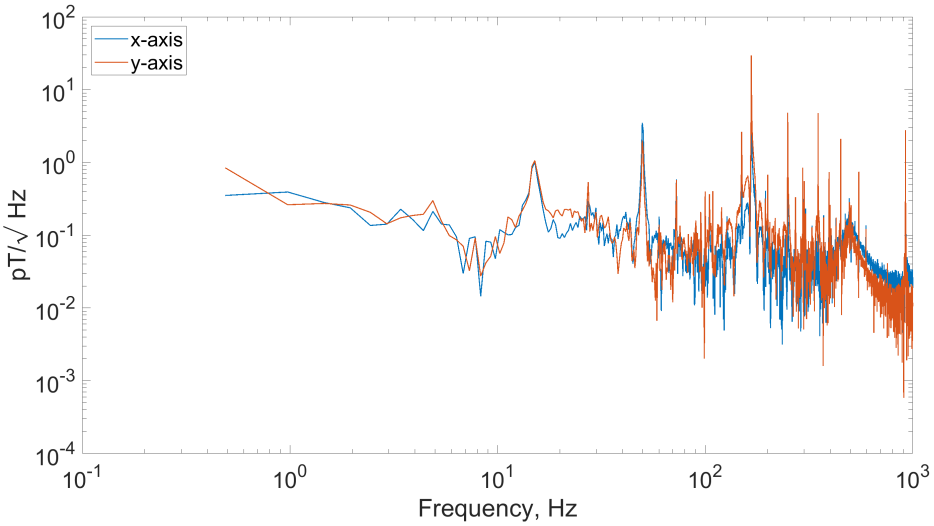


**Figure S4.** Noise floor of the gradiometric magnetometer channels, obtained by taking the power spectral density (PSD) of a single 2 s time trace with an empty sample vial, under experimental conditions and parameters identical to those used in the leaf measurement campaign (Figure S9). Sensors are not responsive above 500 Hz due to an internal cut-off filter; the various background noise sources in the sensitive region below this frequency can be avoided in actual experiment by tuning of the solenoid precession field to an appropriate $^{1}$H Larmor frequency (see Figure S8).

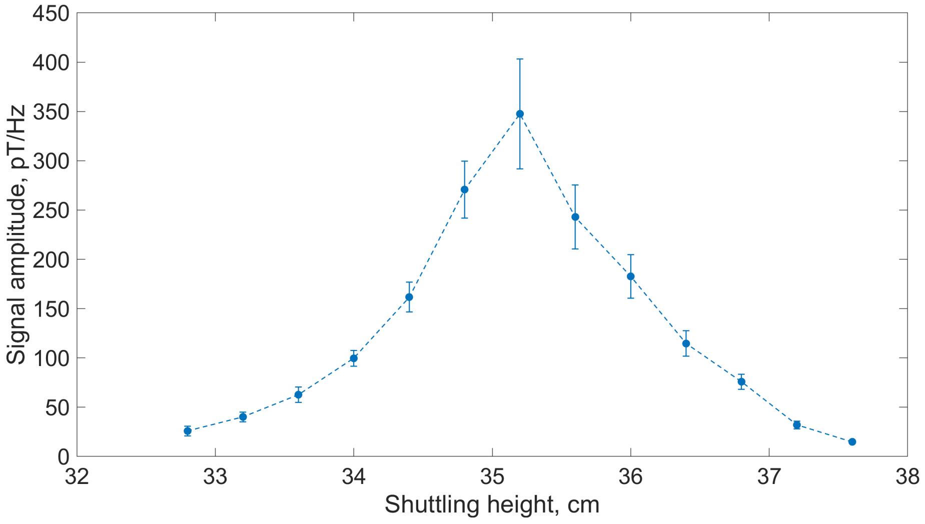


**Figure S5.** Amplitude of the Lorentzian fit to a 10 Hz water calibration signal (average of four scans) as a function of shuttling height above the polarizing magnet. Height of the water column in the sample vial was approximately 2 cm. Error bars have been extracted from fit parameters; connecting dashed lines are a guide to the eye. Experimental settings: 10 s polarization time, 200 µs pulse, 3 s acquisition.

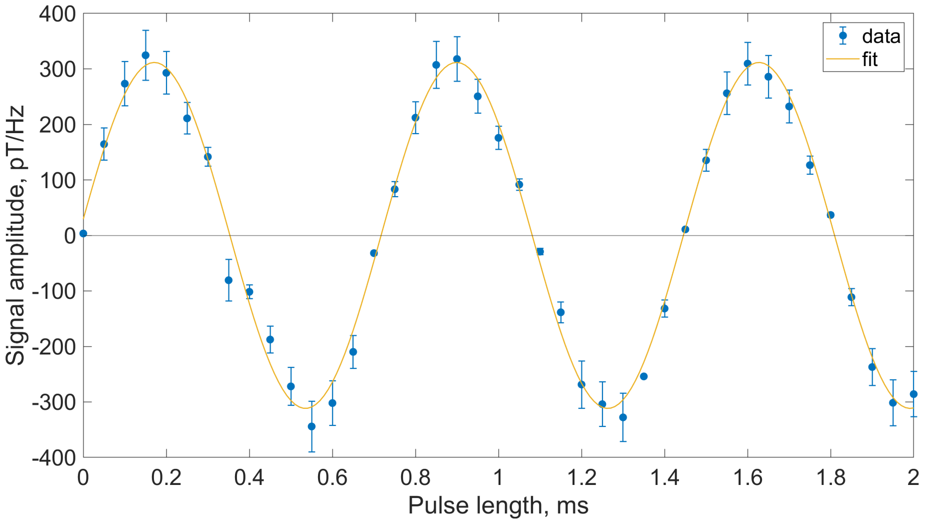


**Figure S6.** Measured nutation curve showing amplitude of the Lorentzian fit to a 10 Hz water calibration signal (average of four scans) as a function of $y$-pulse duration with pulse amplitude 30 µT. A sinusoidal fit is overlaid; error bars have been extracted from Lorentzian fit parameters. Experimental settings: 10 s polarization time, shuttling height 35 cm, 3 s acquisition.

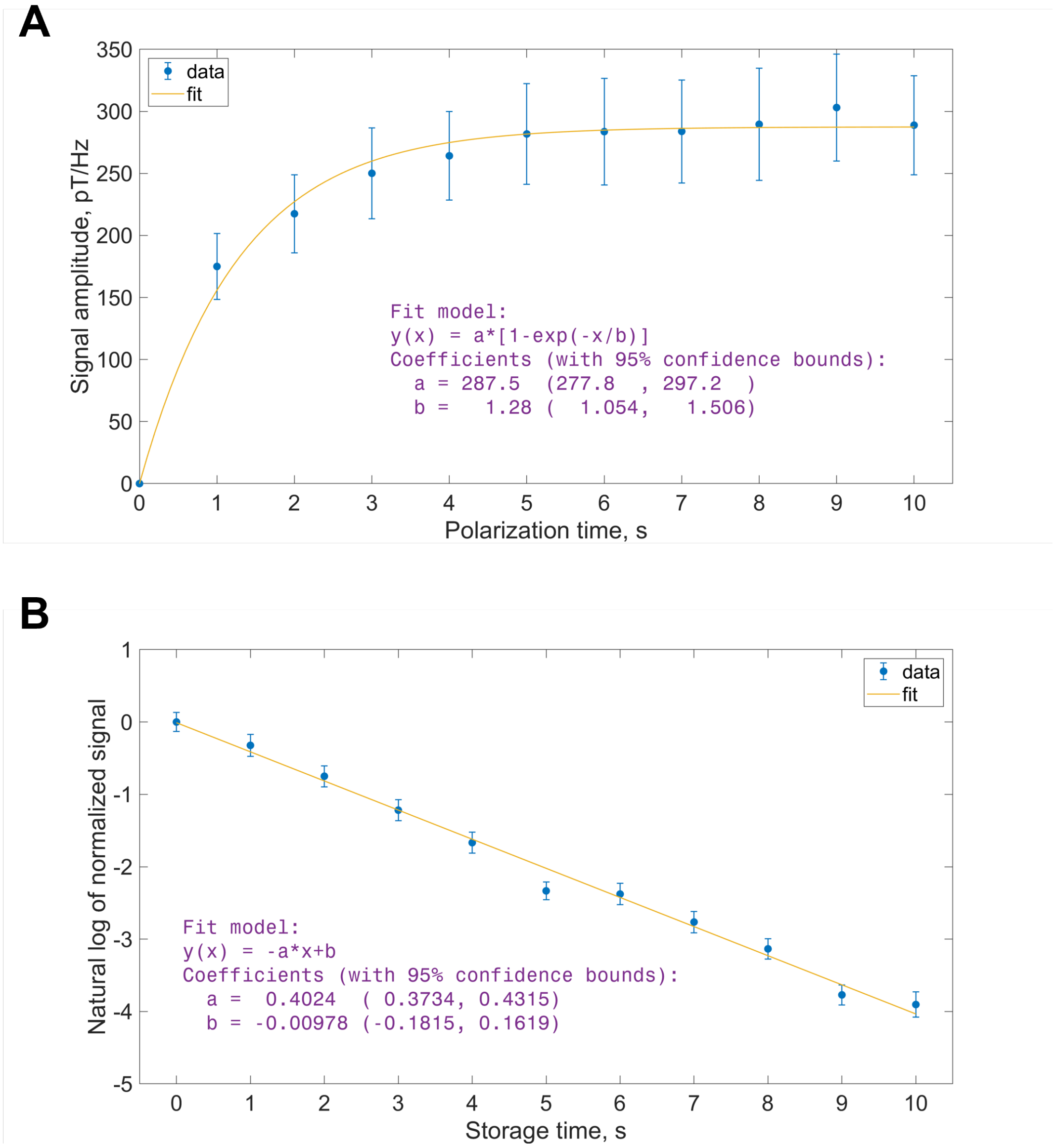


**Figure S7.** Investigation of polarization and storage times using the water calibration sample. **(A)** Amplitude of the Lorentzian fit to a 10 Hz signal (average of three scans) as a function of polarization time $t_\mathrm{p}$ in the Halbach magnet prior to shuttling. Assuming a signal buildup $S(t) = S_\mathrm{p}\left[1 - \exp\left(-t_\mathrm{p}/T_{1,\mathrm{p}}\right)\right]$, where $S_\mathrm{p}$ is the maximal signal corresponding to the maximal achievable spin polarization, we find $T_{1,\mathrm{p}} = 1.28 \pm 0.11$ s. Error bars have been extracted from fit parameters. Experimental settings: shuttling height 35 cm, 200 µs pulse, 3 s acquisition. **(B)** Alternative $T_1$ measurement based on varying the storage time in the solenoid, after shuttling and prior to pulse application. The signal decays exponentially from its maximum value at zero storage time, with a time constant of $T_1$. Taking the inverse slope of the fit, we find $T_1 = 2.49 \pm 0.01$ s. Polarization time was 10 s and four scans were averaged; all other experimental parameters were equivalent to those in the polarization-buildup measurement. Error bars have been extracted from the Lorentzian fit parameters.

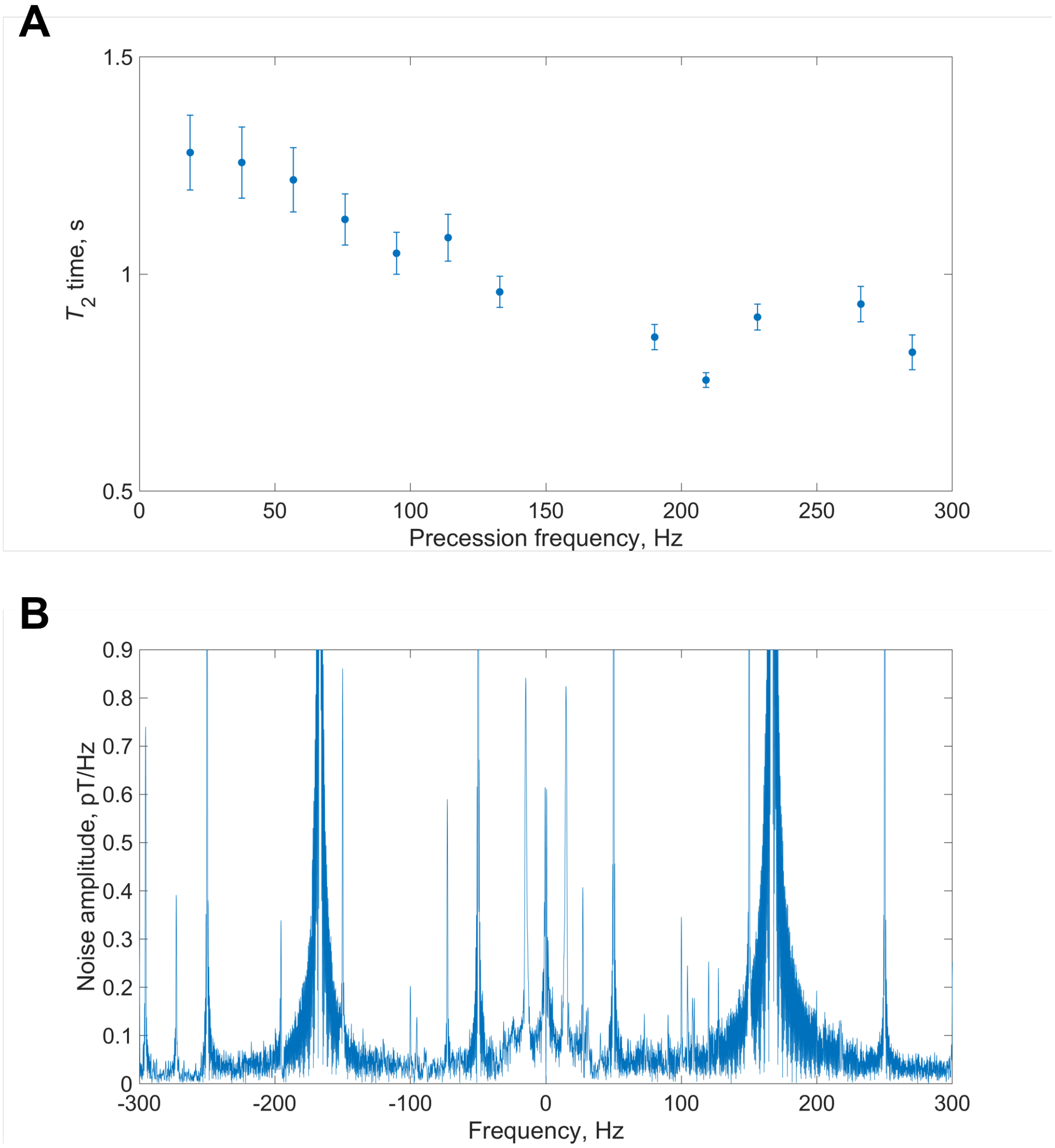


**Figure S8.** Frequency dependence of gradiometric quadrature signal and noise. **(A)** Evolution of measured $T_2$ time as extracted from the width of the Lorentzian fit. Experimental settings: 10 s polarization time, shuttling height 35 cm, 200 µs pulse, 2 s acquisition, four scans. Data points are omitted where the signal peak overlapped with noise peaks. **(B)** Amplitude FFT of the same data set as in Figure S4 (average of four scans). Because signals from leaves are typically three orders of magnitude smaller than those from water calibration signals at a given precession frequency, the relatively noise-free region between 250 and 300 Hz is selected as the best choice for leaf measurements, despite the slight shortening of $T_2$ due to lineshape broadening. Major background noise sources include 50 Hz power-line noise and harmonics thereof, as well as a large unidentified laboratory noise peak at 167 Hz. Given the sampling rate of 2 kHz and associated Nyquist frequency of 1 kHz, smaller noise peaks at $77 \pm 50$ Hz and harmonics are suspected to arise from interference of power-line noise and the 923 Hz internal modulation of the magnetometers.

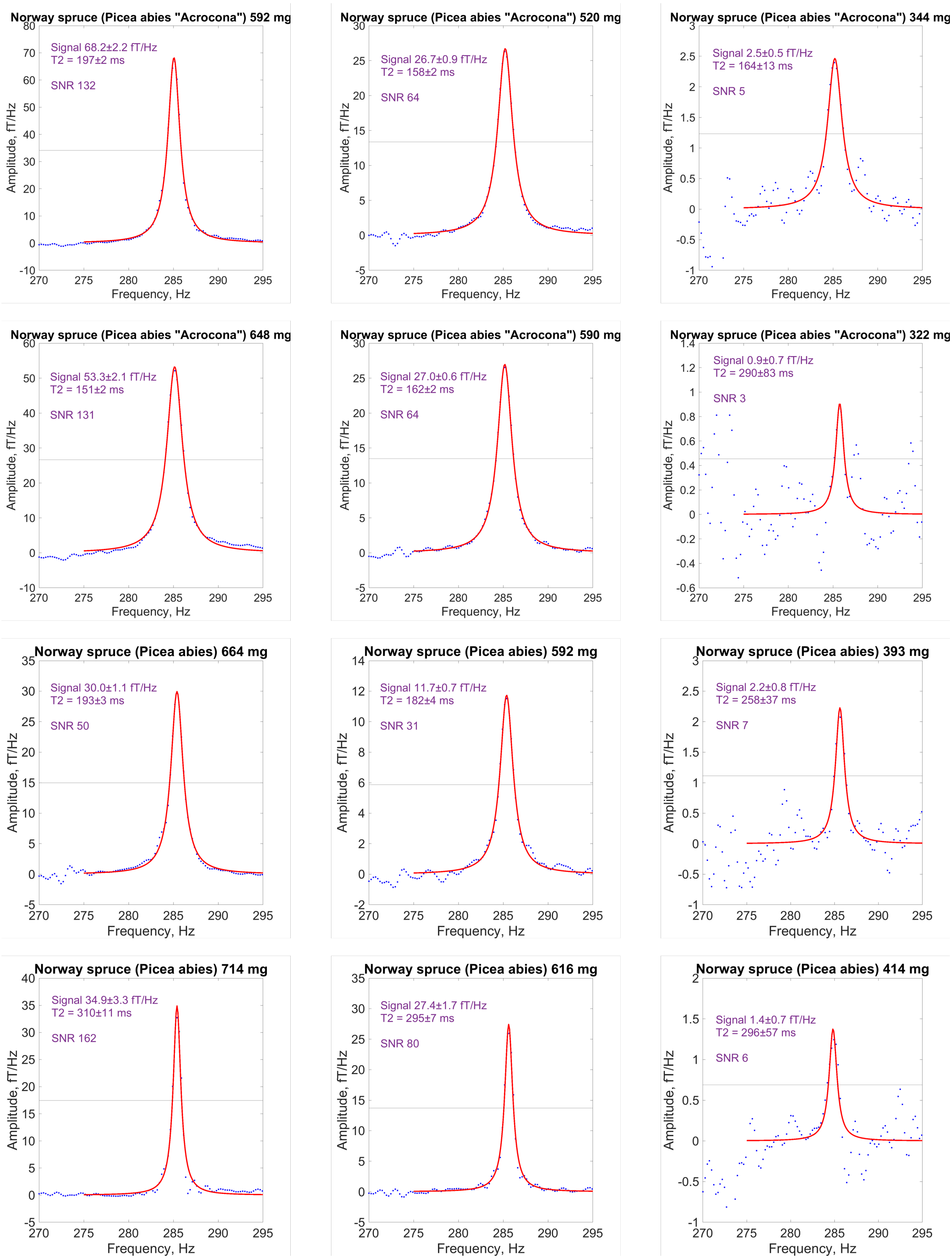


**Figure S9.** Spectra used in the **spruce** dehydration study; each row corresponds to a single sample.

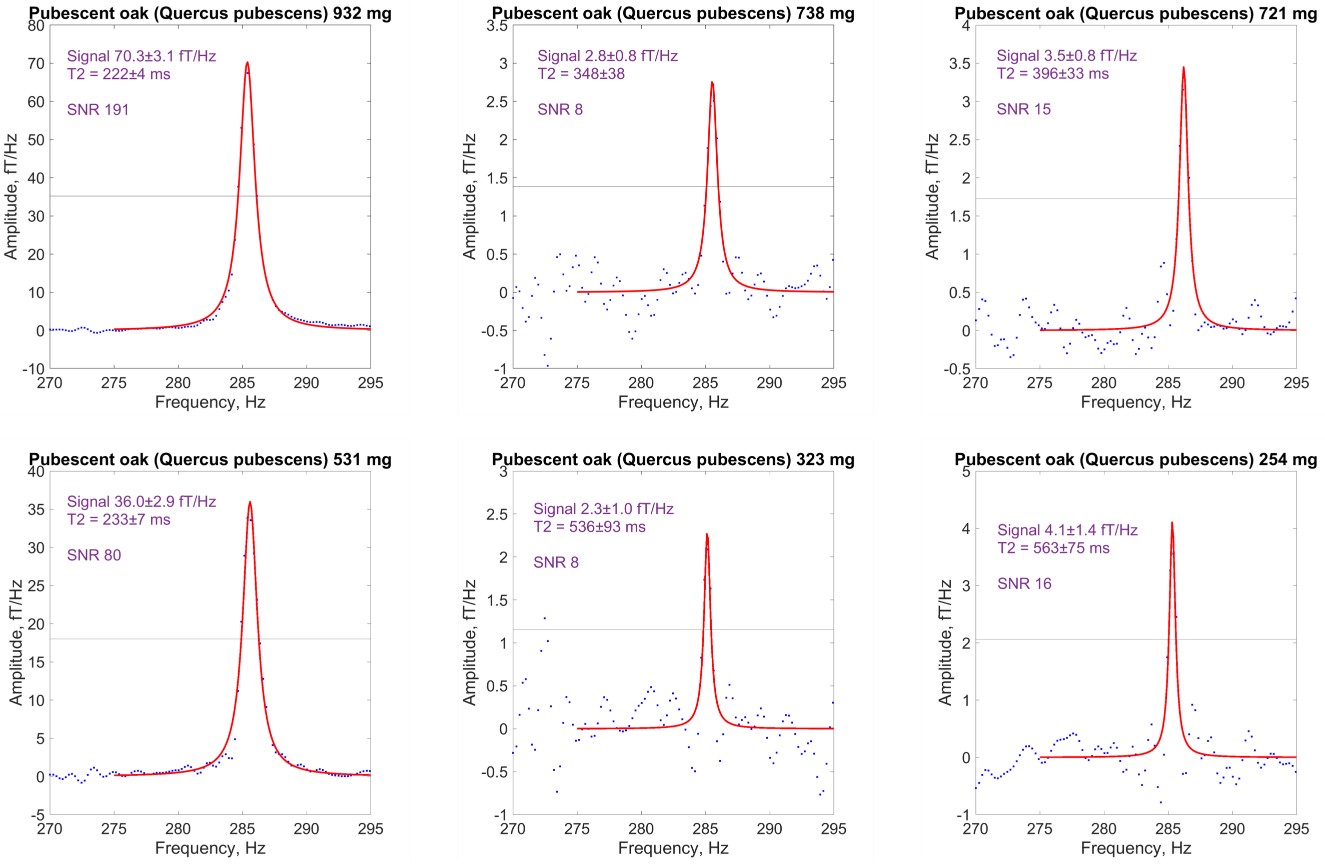


**Figure S10.** Spectra used in the **oak** dehydration study; each row corresponds to the same sample.

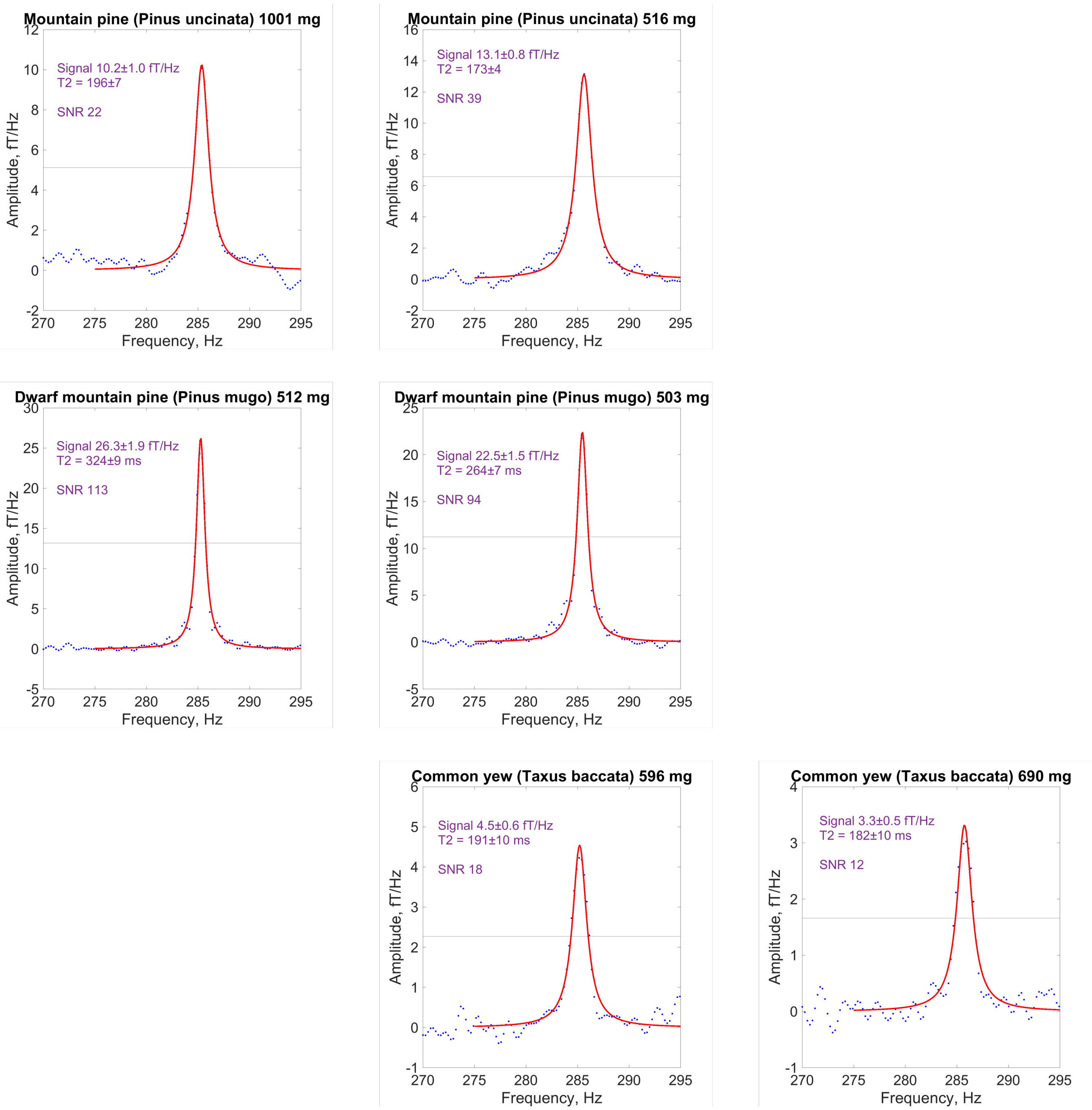


**Figure S11.** Spectra of leaves from evergreen trees including **pine** and **yew**; each plot corresponds to a different sample.

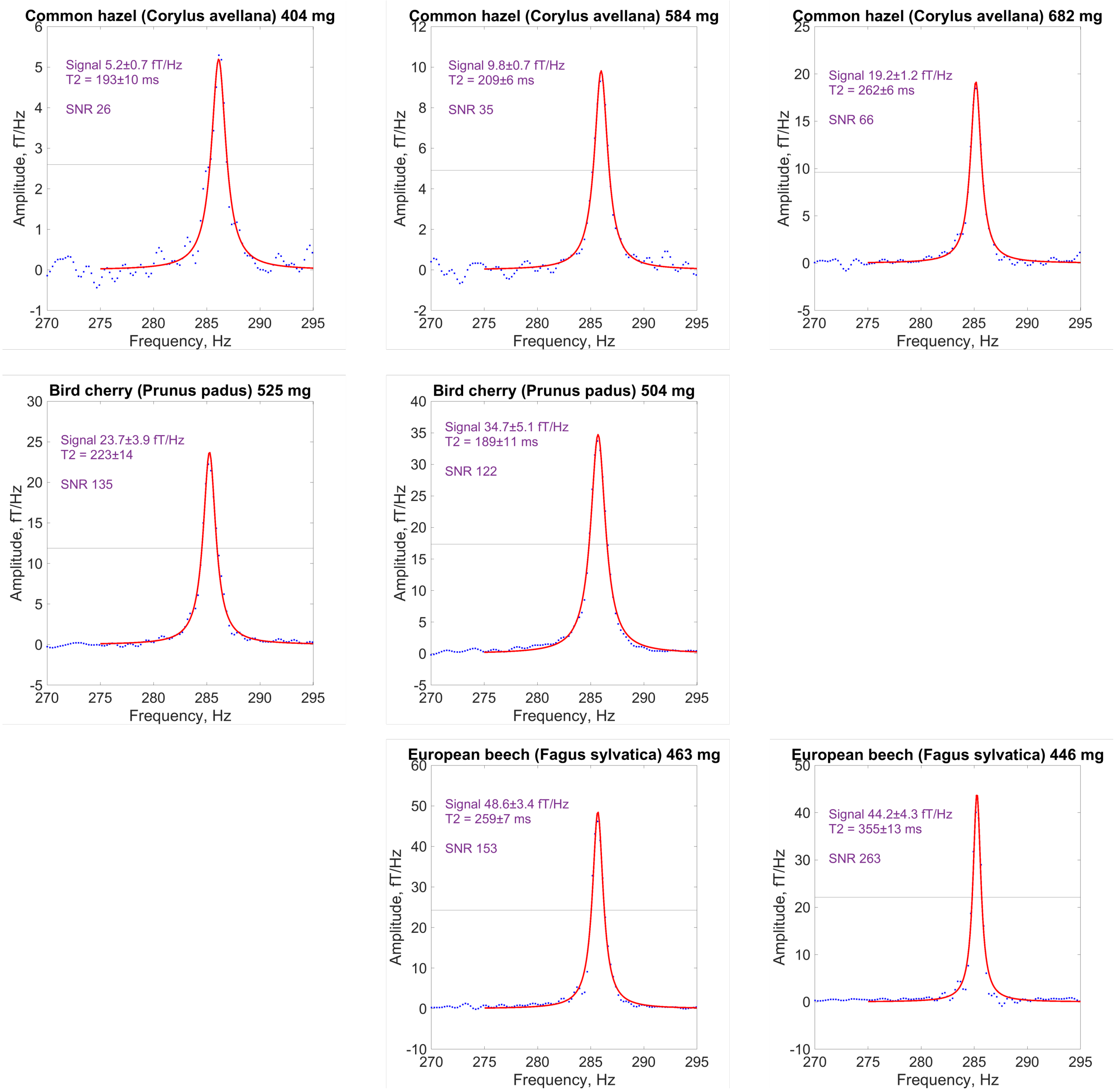


**Figure S12.** Spectra of leaves from deciduous trees including **hazel**, **cherry**, and **beech**; each plot corresponds to a different sample.
